# No phenotypes in *Ttc22* knockout mice

**DOI:** 10.1101/2025.07.20.665798

**Authors:** Jiaoyang Liu, Liankun Gu, Abin You, Hongfan Yuan, Jing Zhou, Wei Tian, Dajun Deng

## Abstract

The *TTC22* gene encodes a protein containing seven tetratricopeptide repeats (TPRs), which mediate protein–protein interactions as chaperones. We previously reported that the level of *TTC22* transcript variant 1 (*TTC22v1*) was downregulated in human colon adenocarcinoma (COAD) and that TTC22 upregulated m6A-mediated *WTAP* and *SNAI1* expression via the TTC22–RPL4 interaction and subsequently promoted COAD metastasis. Thus, a commercially available C57BL/6N mouse model in which the *Ttc22* exon 2&3 encoding the TPR4 (equal to the human TTC22 TPR3) domain was knocked out via CRISPR-Cas9 was used to evaluate the contribution of *Ttc22* to the development of mice and COAD. To determine the status of Ttc22 expression in *Ttc22* knockout mice, we prepared a rabbit polyclonal antibody against the mouse Ttc22 protein, which could bind both wild type Ttc22 and TPR4-deleted Ttc22 proteins overexpressed in HEK293T cells, as determined by Western blot analysis. Ttc22 proteins were detected in colon tissue samples from wild-type mice and *Ttc22* hemi-knockout (*Ttc22 ^-/+^*) mice but not in those from fully *Ttc22* knockout (*Ttc22^-/-^*) mice. Unfortunately, the long-term observation results demonstrated that *Ttc22* knockout (KO, including *Ttc22*^-/+^ or *Ttc22*^-/-^) did not affect the body weight, development, fertility, or spontaneous tumor incidence of male or female mice. No differences in the incidence of AOM/DSS-induced COAD were observed between these mouse groups, although *Ttc22* deletion partially resulted in the upregulation of genes related to the host adaptive response to inflammation in the colon mucosa. In conclusion, no phenotypes were induced by *Ttc22* inactivation in C57BL/6N mice.

**Highlights:**

- Knockout of *Ttc22* exons 2 and 3 completely disrupted gene expression.
- Knockout of *Ttc22* does not affect the development of C57BL mice.
- Knockout of *Ttc22* does not affect the induction of mouse colon cancer by AOM/DSS.

## INTRODUCTION

The human TTC22 protein contains seven tetratricopeptide repeats (TPRs), which mediate protein-protein interactions and are involved in multiple biological processes, including the movements of cilia and flagella and the migration of human cells [1-5]. A pilot study revealed that the *TTC22* gene was among the top ten downregulated genes in the fetuses of obese women compared with lean women in the second trimester and that low *TTC22* expression in mothers may be related to fetal neurodevelopment and metabolism [6]. Differentially methylated regions of the *TTC22* gene are associated with hand grip strength [7]. DNA sequencing and transcriptome data analyses in 3,203 subjects suggested that low-frequency variants of *TTC22* were associated with low-density lipoprotein (LDL) cholesterol [8]. These phenomena imply that TTC22 may be involved in physiological metabolism. However, the causality between TTC22 alterations and metabolism is not known.

According to data from the Human Protein Atlas and The Cancer Genome Atlas (TCGA), the *TTC22* gene is comprehensively expressed in epithelial cells in the normal gut mucosa at the middle level and is extensively downregulated in gastrointestinal cancer tissues [9,10]. We initially reported that *TTC22* variant 1 (*TTC22v1*) mRNA is targeted by *miR663a*, an anti-inflammatory miRNA in the colon [11]. *TTC22v1* mRNA expression is significantly lower in human colon adenocarcinoma (COAD) than in paired normal tissue samples, and high *TTC22* expression increases the risk of colon cancer metastasis. Our further study revealed that TTC22 increased m6A-mediated *WTAP* and *SNAI1* expression via TTC22-RPL4 binding and subsequently promoted COAD metastasis *in vitro* and *in vivo* [12], suggesting a potential role for TTC22 in the development and progression of COAD.

Therefore, we further studied the effects of Ttc22 on the development of mice and their susceptibility to the chemical carcinogen AOM/DSS via a genetically established and commercially available C57BL gene knockout mouse model in which *Ttc22* exons 2 and 3, which encode the TPR4 domain of the mouse Ttc22 protein, were knocked out via CRISPR–Cas9. While qRT-PCR and Western blot analyses confirmed the complete absence of *Ttc22* mRNA and protein in the colon tissues of mice with homozygous deletion of two *Ttc22* alleles (*Ttc22*^*-/-*^), unexpectedly, no differences in the development, growth, fertility, or AOM/DSS susceptibility of the mice were induced by *Ttc22* knockout. Transcriptomic analysis of colon tissues revealed significant upregulation of genes involved in protein folding and stress response in *Ttc22*^*-/-*^ mice, indicating that loss of *Ttc22* induces a broad adaptive response that may compensate for its absence. Our findings demonstrate that Ttc22 is dispensable under physiological conditions because of gene redundancy or genetic robustness, highlighting the importance of compensatory mechanisms in gene function studies.

## MATERIALS AND METHODS

### *Ttc22* exon 2&3 knockout mouse model

Genetically established *Ttc22* knockout (Ttc22-KO) C57BL/6N mice (strain S-KO-06335) were purchased from Cyagen (Cyagen Bio. Co., Guangzhou, China). Ttc22 knockout mice were obtained via high-throughput electroporation of fertilized eggs. The CRISPR-Cas9 system and two gRNAs (gRNA#1: 5’-ggaaagugggcauaacgccuggg-3’ and gRNA#2: 5’-uuccagagagauagcgaccgagg-3’) targeted to Ttc22 introns 1&3 were used to delete a 1610 bp fragment including *Ttc22* exons 2&3 encoding the Ttc22 TPR4 domain (Figure 1A). After sexual maturity, the sperm were collected for cryopreservation. Through hybridization with female wild-type C57BL/6N mice, first-generation Ttc22^+/-^ mice were obtained and used to generate male and female mice with different *Ttc22* genotypes (including Ttc22^+/+^, Ttc22^+/-^, and Ttc22^-/-^), as determined by PCR sequencing via the tail DNA template and two forward primers (OuterF: 5’-tcacaacctactcaagatccagc-3’ and InnerF: 5’-caactgtagggtatagcagtccc-3’) and one common reverse primer (OuterR: 5’-ttattgcctccttggtgagttg-3’). A mouse tissue direct PCR kit (YEASEN, Shanghai, China) was used to amplify *Ttc22*. Both male and female *Ttc22*^*+/+*^ and *Ttc22*^*-/-*^ mice were maintained as seed mice in an animal facility under SPF conditions at the Peking University Cancer Hospital & Institute. The seed mice were hybridized to generate experimental mouse stocks with different *Ttc22* genotypes. The body weight of each mouse was measured once a week from 4–31 weeks of age. The animal experiments were approved by the Biomedical Ethical Committee of Peking University Cancer Hospital & Institute.

**Figure 1.**
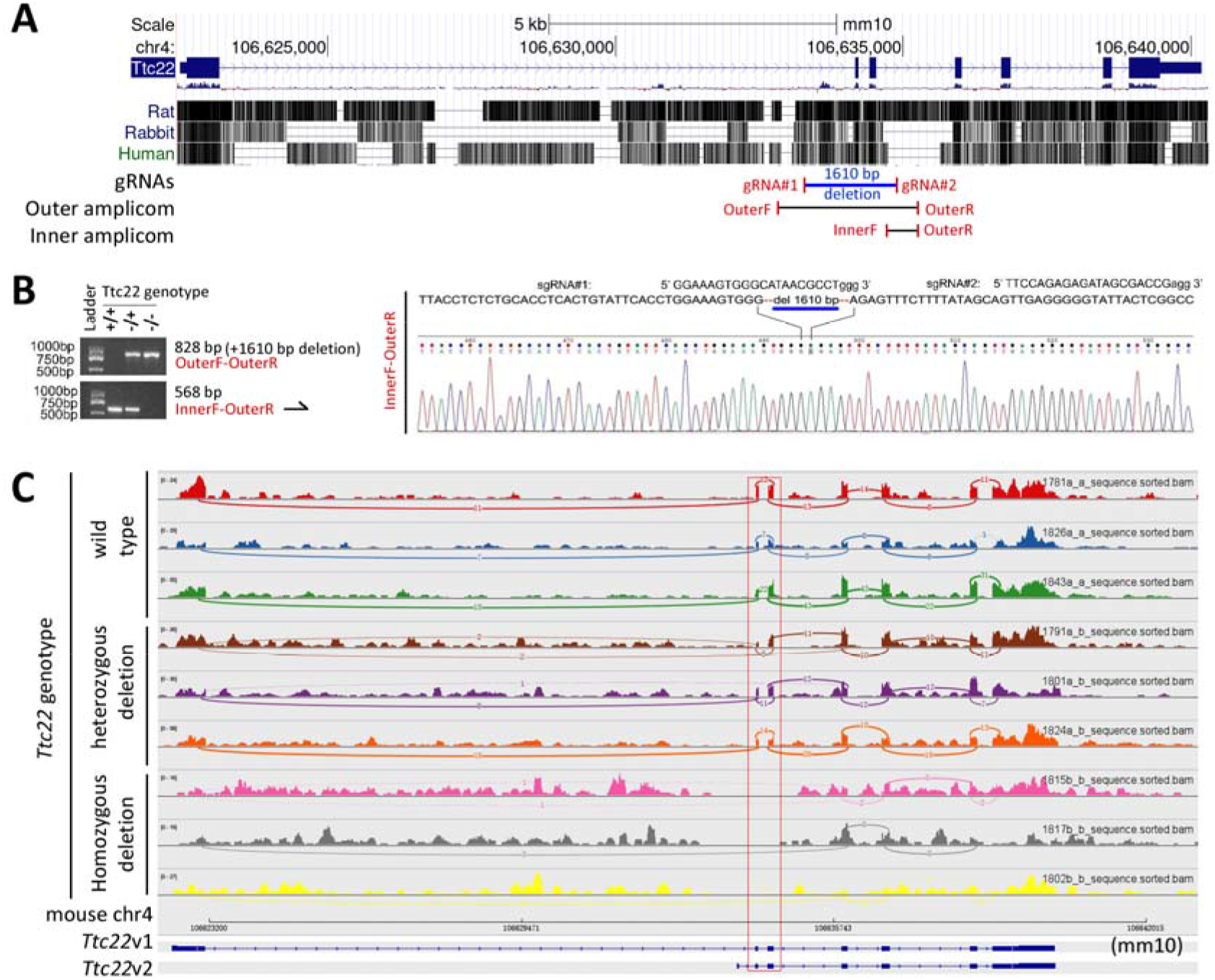
Characterization of the genotypes of the *Ttc22* gene in C57BL/6N mice via CRISPR-Cas9. (A) Locations of the deleted fragment (1610 bp), amplicons, gRNAs, and primers matched to the *Ttc22* gene. The conservation status of *Ttc22* in placental mammals, including rats, rabbits, and humans, is displayed below. (B) Characterization of *Ttc22* genotypes, as determined by PCR and sequencing. The wild-type *Ttc22* fragment is too long (2438 bp) to be amplified via regular PCR via the OuterF and OuterR primer sets. (C) IGV view of *Ttc22* transcripts in RNA samples from the colons of three male mice with different *Ttc22* genotypes, namely, *Ttc22* wild-type (*Ttc22*^*+/+*^), heterozygously deleted (*Ttc22*^*-/+*^), and homozygously deleted (*Ttc22*^*-/-;*^), strains, as determined by RNA sequencing. The exon 2&3 region, which is shared by both *Ttc22* variants 1 and 2 and is knocked out by CRISPR-Cas9, is highlighted by a red rectangle.

### High-fat diet experiment

Four-week-old female C57BL/6N mice were consistently fed a high-fat diet, in which 45% of the calories were derived from fat (4.73 kcal/g H10045, Huafukang Bio. Co., Beijing, China). Additionally, a control group of four-week-old female mice was fed a regular diet. The mice were housed in a temperature-controlled room with a 12-h light/dark cycle. All the mice had free access to food and water [13].

### Fertility test

Mating tests were carried out on 6–8-week-old mice. One male Ttc22^+/+^ mouse was mated with 1 Ttc22^+/+^ female mouse as the control group, and 1 male Ttc22^-/-^ mouse was mated with 1 Ttc22^-/-^ female mouse as the Ttc22-KO group for the first pregnancy. At least three mates were included in each group. The date and number of pups were recorded at birth in each cage.

### Induction of COAD by AOM/DSS

Five male mice aged 6–8 weeks were included in each group within the same cage. The mice in the three groups, *Ttc22*^*+/+*^, *Ttc22*^*+/-*^, and *Ttc22*^*-/-*^ were intraperitoneally administered 10 mg/kg b.w. azoxymethane (AOM) solution (1 mg/mL, freshly prepared with saline; Cat. #A5486 Sigma–Aldrich; Merck KGaA, Darmstadt, Germany) on the 1st experimental day. Beginning on the 2nd experimental day, the mice were administered dextran sodium sulfate (DSS; Cat. #24867; Sigma) in the drinking water containing 2.5% DSS (2.5 g/100 mL) for 7 continuous days. The mice were subsequently allowed to recover for three weeks (drinking water not containing DSS). The combined AOM/DSS treatment cycle was repeated two more times [14]. From the 24th to 27th experimental weeks, the mice were sacrificed, the colon was collected, and the length of the colon was measured. The intestine was opened, and the colon mucosa was carefully examined and photodocumented. Evidence of gross pathologic changes was recorded (e.g., ulceration). Afterward, the colon tissues were fixed with 10% formalin, rolled Swiss, embedded in paraffin, and sectioned for pathological analysis. RNA or protein was extracted from a portion of fresh colon tissue with TRIzol or cell lysis buffer. The remaining tissue was stored at -80°C for subsequent studies. The roll was further fixed in 10% buffered formalin overnight at room temperature and used to prepare tissue sections (4 μm). Pathology was performed by senior pathologists.

### RNA sequencing

Total RNA was extracted from the colon tissues of three male mice with different *Ttc22* genotypes via an RNA extraction kit. RNA sequencing was performed by Cloud-Seq Biotech (Shanghai, China). Briefly, total RNA was removed from the rRNAs with an NEBNext rRNA depletion kit (Biolabs, Inc., Massachusetts, USA), which was subsequently used to construct RNA libraries via the NEBNext Ultra II directional RNA library prep kit (Biolabs). Library sequencing was performed on an Illumina NovaSeq 6000 instrument with 150 bp paired-end reads. Paired-end reads were subsequently harvested. The high-quality clean reads were aligned to the reference genome with HISAT2 software (v2.0.4), guided by the Ensembl gtf gene annotation file, and the gene level FPKM was obtained as the expression profile of the mRNAs. Fold changes and p values were calculated on the basis of the FPKM values, and the differentially expressed mRNAs were identified on the basis of |log(FC)|>=1.0 and a p value<= 0.05. GO and pathway enrichment analyses were performed on the differentially expressed mRNAs. The raw data were deposited under the Gene Expression Omnibus accession number GSE275363.

### m6A dot blot analysis

m6A dot blot analysis was performed as we previously reported [12].

### Quantification of m6A abundance via m6A-specific ELISA

The total RNA m6A level was quantified with an EpiQuik m6A RNA Methylation Quantification Kit (P-9005; Epigentek Group, NY, USA) according to the manufacturer’s instructions [15,16].

### LC–MS/MS measurement of the m6A/rA ratio

Liquid-MS/MS quantification of m6A was performed by Cloudseq Biotech, Inc. (Shanghai, China), as described previously [12].

### Full-length Ttc22, ΔTPR1, and ΔTPR4 expression vectors

The pCMV6-Flag full-length Ttc22 expression vector was purchased from the MiaoLing Plasmid Platform (Wuhan, China) [12]. The ΔTPR4 vector not containing 568-738 bp and the ΔTPR1 vector not containing 4-297 bp were derived from the full-length Ttc22 vector and confirmed by Sanger sequencing (Figure S1-S3). These vectors were used to transiently transfect human colon cancer HEK293T cells (ATCC; Manassas, VA, USA).

### Western blot analysis

Cells were harvested 48 hrs after transfection and in 1× cell lysis buffer [50 mmol/L Tris-HCl (pH 6.8), 100 mmol/L DTT, and 2% SDS. 0.1% bromophenol blue and 10% glycerol] was used to extract total cellular proteins. Protein electrophoresis was performed using 10% SDS-PAGE gels. The membranes were incubated with the following primary antibodies at 4°C overnight: a mouse anti-DYKDDDDK Flag monoclonal antibody (1:3000; cat. no. 66008-4-Ig, Proteintech Group, Inc. Wuhan, China), a mouse anti-Gapdh monoclonal antibody (1:10000; cat. no. 60004-1-Ig; Proteintech Group, Inc., Wuhan, China), and a rabbit anti-His-TTC22 (1-569 aa) polyclonal antibody previously raised and characterized in our own laboratory [12]. The membranes were subsequently incubated with horseradish peroxidase (HRP)-conjugated anti-rabbit or anti-mouse secondary antibodies (1:5000; cat.no. A0208 or A0216, Beyotime Technology, Shanghai, China) at room temperature for 50 min.

Mouse colon tissue was collected and homogenized in a 1.5 mL microcentrifuge tube. Afterward, 1 mL of lysis buffer was added per 50 mg of tissue. The remaining steps were carried out as described above for the cell samples.

### Statistical analysis

Statistical analysis was carried out via SPSS 22. 0 software (SPSS Inc., Chicago IL, USA). The data are presented as the means ± SDs of at least three independent experiments or as the medians (25th–75th percentiles). Statistical analysis methods included Student’s t test, one-way analysis of variance (ANOVA), and two-way ANOVA. The exact statistical methods are described in detail for each experiment in the Results section. All tests were two-sided. A p value <0.05 was considered to indicate statistical significance.

## RESULTS

### Characterization of *Ttc22* genotypes

To evaluate the effects of the *Ttc22* gene on the development of mice and COAD, a commercially available heterozygous Ttc22-KO C57BL/6N mouse model was used to generate mice with various *Ttc22* genotypes, including homozygous and heterozygous Ttc22-KO (*Ttc22*^*-/-*^ and *Ttc22*^*+/-)*^ and its wild-type control (*Ttc22*^*+/+*^) mice, as described in the Methods section. The sequencing results indicated that a 1610 bp fragment, including *Ttc22* exons 2 and 3, which encode the Ttc22 TPR4 domain (57 aa), was deleted via CRISPR-Cas9 and two gRNAs (Figure 1A and 1B). RNA sequencing confirmed the sequencing results (Figure 1C).

To determine the expression level of the Ttc22 protein in the colons, we performed Western blot analysis using a rabbit polyclonal antibody against human TTC22 previously created at our laboratory [12]. We initially investigated whether the TTC22 antibody might also bind mouse Ttc22 and found that the TTC22 antibody efficiently bound both wild-type Flag-Ttc22 and its ΔTPR1 and ΔTPR4 variants (Figure 2A and Figure S1-S3) overexpressed in HEK293T cells by Western blotting (Figure 2B). These results were consistent with those obtained with the anti-Flag antibody. However, Ttc22 was detected only in the colon tissues from *Ttc22*^*+/+*^ and *Ttc22*^*+/-*^ male mice, and no Ttc22 was detected in the colon tissues from *Ttc22*^*-/-*^ mice (Figure 2C), indicating complete loss of Ttc22 expression in the colon. We further checked the knockout status of the *Ttc22* alleles in the colon tissues from these mice. To simultaneously amplify both wild-type *Ttc22* and its exon 2&3-deleted variant by the same PCR, we pooled three primers (OuterF, *Ttc22* InnerF, and OuterR) together. As expected, the 828 bp PCR products were amplified only by OuterF and OuterR primers from the genetic DNA of *Ttc22*^*-/-*^ and *Ttc22^+/-^* mice and were not amplified from the genetic DNA of *Ttc22*^*+/+*^ mice (1610 bp + 828 bp = 2438 bp; too long to be amplified). In contrast, the 568 bp PCR products were amplified only by the InnerF and OuterR primers in *Ttc22*^*+/+*^ and *Ttc22*^*+/-*^ male mice and not in *Ttc22*^*-/-*^ mice (Figure 1D). Collectively, these results indicate that only *Ttc22* exons 2 and 3 are knocked out in these mice, which leads to complete loss of Ttc22 expression.

**Figure 2.**
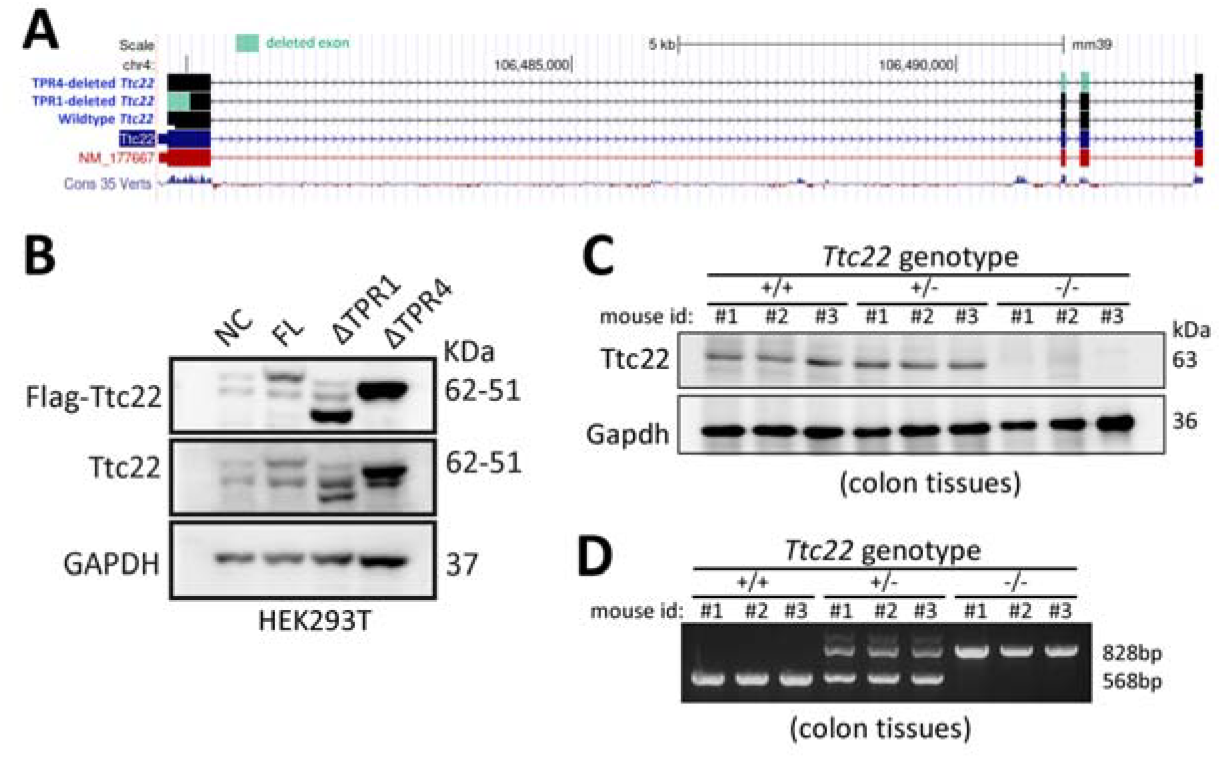
Characterization of the knockout status of the *Ttc22* gene in the mouse colon. (A) Illustration of wild-type *Ttc22* and its TPR1- or TPR4-deleted (ΔTPR1 or ΔTPR4) expression vectors. (B) Western blot images of the full-length (FL)/wild-type Ttc22 protein and its ΔTPR1 or ΔTPR4 variant overexpressed in HEK293T cells. (C) The abundance of the Ttc22 protein in mouse colon tissues from three groups of mice, as determined by Western blotting. (D) Characterization of *Ttc22* genotypes in mouse colon tissues, as determined by PCR via a pooled primer set, including OuterF, InnerF, and OuterR.

### No morphologic alterations in heterozygous and homozygous Ttc22-KO mice

Compared with wild-type C57BL/6N mice, those with different *Ttc22* knockout states did not differ in development, growth, appearance (Figure 3), body weight (Figure 4A and 4B), or fertility (Figure 5).

**Figure 3.**
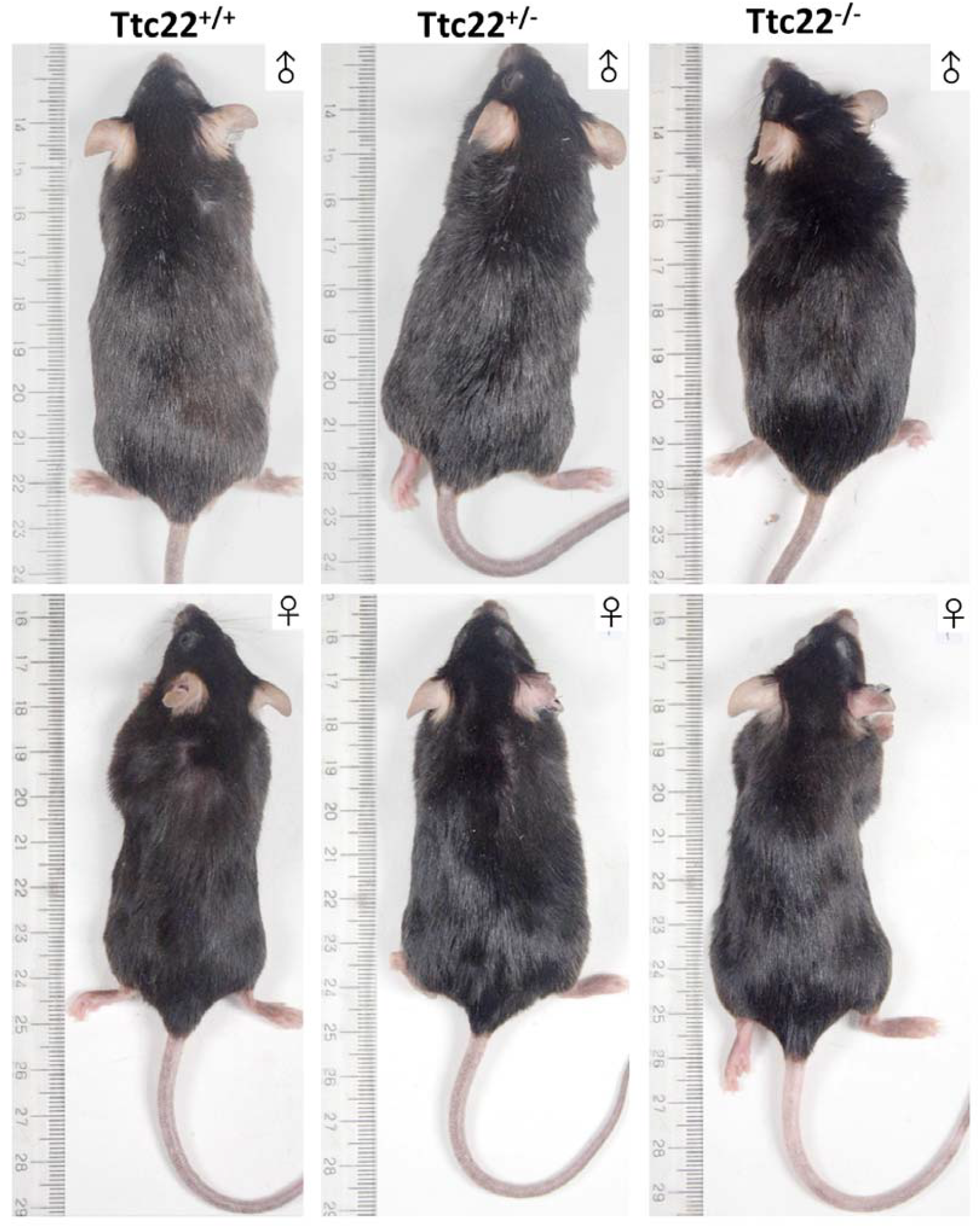
The appearance of C57BL/6N male and female C57BL/6N mice with different *Ttc22* genotypes at approximately 66 and 77 weeks of age

**Figure 4.**
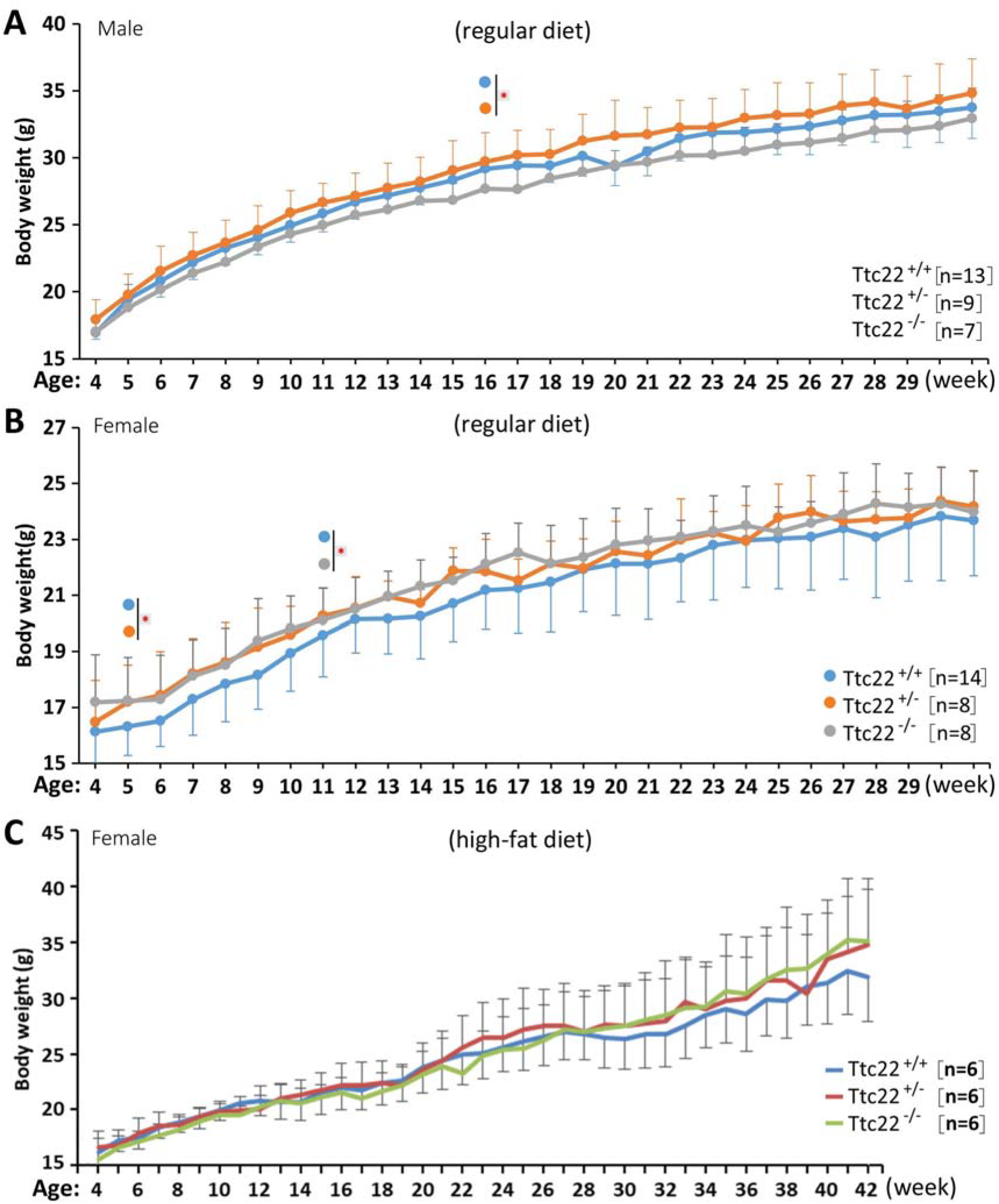
Body weight curves of C57BL/6N mice in different *Ttc22* genotype groups fed a regular or high-fat diet. (A and B) Body weight curves for male and female mice fed a regular diet. *p<0.05 between groups was observed only on three days (the 16th week for males and the 5th and 11th weeks for females) during 31 experimental weeks, according to Student’s t test. (C) Body weight curves for female mice fed a high-fat diet. The dots and bars represent the mean values and standard deviations (SDs) of the body weights of the mice. *p<0.05 according to Student’s t test.

**Figure 5.**
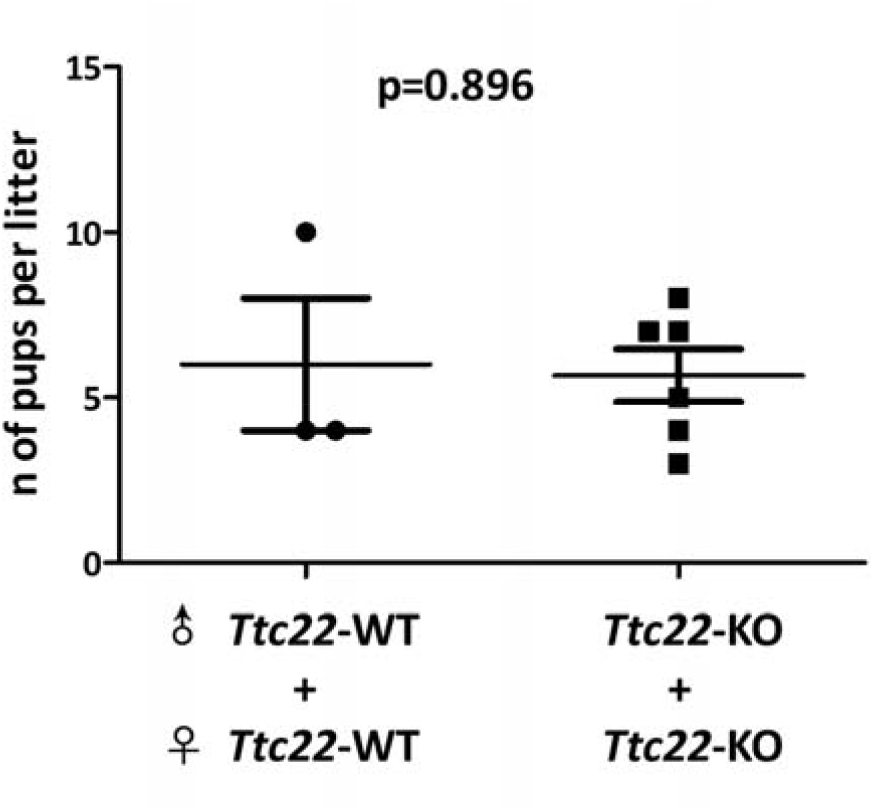
Fertility comparison between *Ttc22*^*+/+*^ and *Ttc22*^*-/-*^ mice. Mating pairs were monitored for litter production during the first pregnancy. Three pairs of wild-type (*Ttc22*^*+/+*^) mice and six pairs of homozygous knockout (*Ttc22*^−*/*−^) mice were established. Each symbol represents a pair. The data are presented as the mean ± SD.

Considering that *TTC22* is mostly downregulated in the fetuses of obese women and that low-frequency variants of TTC22 are associated with low-density lipoprotein (LDL) cholesterol in the human body [1,3], we further compared the body weights of female *Ttc22* mice fed a high-fat diet. However, no differences in the body weights of C57BL/6N mice in different *Ttc22* genotype groups were observed (Figure 4C).

No spontaneous tumors were observed in these mice up to 40 weeks of age. As expected, spontaneous tumors, including lymphoma, sebaceous carcinoma, and basal carcinoma, occurred in some old mice. However, no differences in the incidence of spontaneous tumors were detected between these *Ttc22* genotype groups (Table 1).

**Table 1.** Incidence and latency period of spontaneous tumors* in C57BL/6N mice with different *Ttc22* genotypes.

| Genotype | Male |  |  | Female |  |  |
| --- | --- | --- | --- | --- | --- | --- |
|  | *n* | Mice with tumor (%) | Range of tumor latent weeks | *n* | Mice with tumor (%) | Range of tumor latent weeks |
| *Ttc22*<sup>+/+</sup> | 17 | 3 (17.6) | 45, 55, 77 | 12 | 0 | - |
| *Ttc22*<sup>+/-</sup> | 14 | 2 (14.3) | 57, 60 | 19 | 4 (21.1) | 57, 60, 65, 75 |
| *Ttc22*<sup>-/-</sup> | 27 | 2 (7.4) | 54, 54 | 39 | 6 (15.4) | 41, 55, 55,55, 59, 60 |
| (sum) | 58 | 7 (12.1) |  | 60 | 10 (16.7) |  |

\* including lymphoma, sebaceous carcinoma, and basal carcinoma

### No differences in AOM/DSS-induced COAD between *Ttc22*^*+/+*^, *Ttc22*^*+/-*^, and *Ttc22*^*-/-*^ mice

Because *TTC22* is involved in COAD development and metastasis [5,6], we wondered whether Ttc22-KO might affect the development of COAD induced by three cycles of AOM/DSS treatment in male mice. By the end of the 24th or 27th experimental week, well-differentiated or moderately differentiated adenocarcinomas were induced in the colons of all AOM/DSS-treated mice but not in those of the 0.9% NaCl control mice (Figure 6). The AOM/DSS-induced tumor burden in *Ttc22*^*+/+*^ mice was heterogeneous, ranging from multiple small nodules to a large tumor (d =1 cm). Similar phenomena were also observed in *TTC22*^*+/-*^ mice, with one animal harboring a very large tumor (d=1.8 cm). Two of the five AOM/DSS-treated *Ttc22*^*-/-*^ mice experienced fatalities during the experimental period. However, no differences in the incidence or severity of colorectal tumors were observed between these mouse groups.

**Figure 6.**
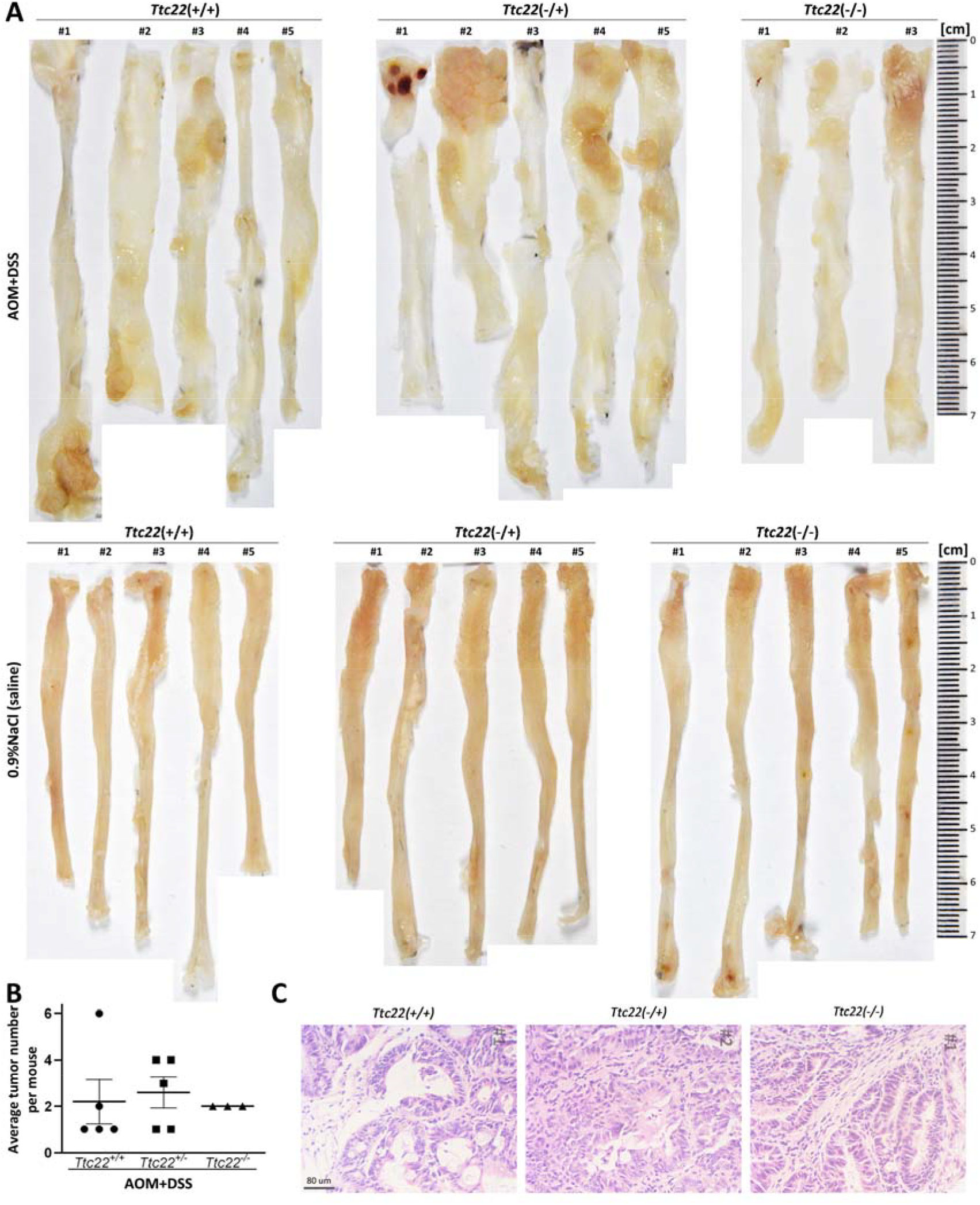
Induction of colon tumors by AOM/DSS in male C57BL mice with different *Ttc22* genotypes. (A) Overview of tumors in the prefixed colon; (B) Scatter plot showing the average tumor number in the colon per mouse; each dot represents one mouse. The data are presented as the mean ± SD. (C) Histological images of representative tumors induced by AOM and DSS.

### Changes in genome-wide adaptive gene expression in the colon of Ttc22-KO mice

We performed RNA sequencing on total RNA extracted from the colons of three 58-week-old male mice with different *Ttc22* genotypes. A total of 528 genes were differentially expressed (|log(FC)|≥1 and p≤0.05) upon *Ttc22* knockout, comprising 303 upregulated genes and 225 downregulated genes. The results of the GO and pathway analyses revealed that *Ttc22* knockout resulted in the upregulation of genes related to infection by microorganisms (Figure 7).

**Figure 7.**
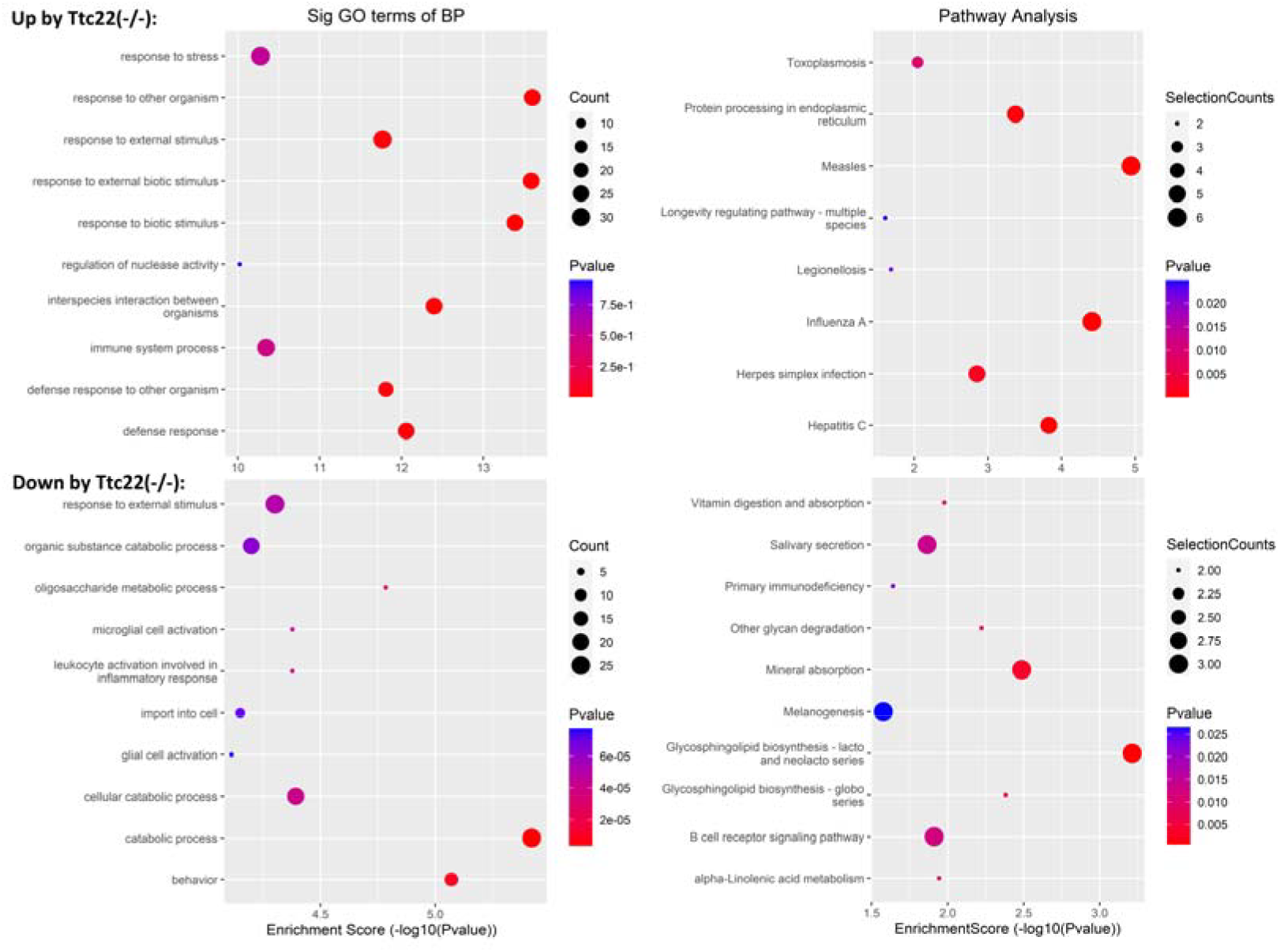
Enrichment scores for genes upregulated and downregulated by homozygous *Ttc22* knockout in the colon of 57-week-old male mice

We previously reported that TTC22 increases total RNA m6A modification through RPL4-mediated WTAP upregulation [12]. Interestingly, no significant difference in the total RNA m6A level was observed in the colons between these groups of mice with different Ttc22 genotypes (Figure 8).

**Figure 8.**
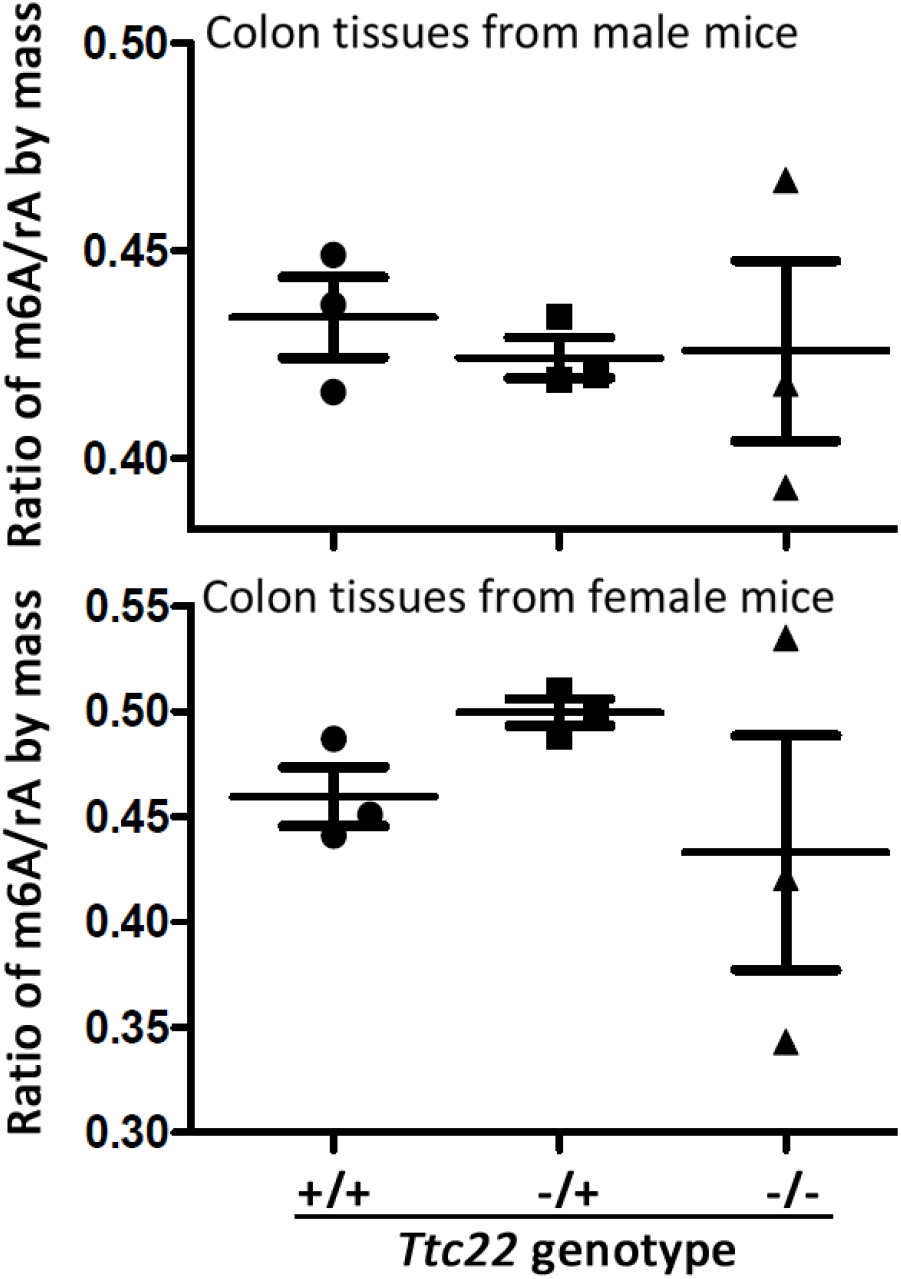
Effect of Ttc22 inactivation on total RNA m6A modification in the colons of mice. The ratio of total m6A to total adenosine in RNA (m6A/rA) in the colon tissues of mice with various *Ttc22* genotypes, as determined by mass.

The above results imply that Ttc22 may contribute to host defense responses against various infections and colon cancer development and that *Ttc22* knockout induces a broad adaptive response that may compensate for its absence.

## DISCUSSION

Considering that *TTC22* mediates human COAD metastasis [11,12], we studied the influence of Ttc22 knockout on the development of COAD in C57BL/6N mice. Unfortunately, no significant differences in the development or fertility of mice and AOM/DSS-induced COAD were observed between wild-type and Ttc22-KO C57BL/6N mice, although deletion of the Ttc22 TPR4 domain fully disrupted *Ttc22* expression in mice and the mechanism underlying this disruption is unknown.

The AOM/DSS-mediated COAD model is a conventional and efficient model for studying inflammation-driven colon carcinogenesis in a controlled experimental timeframe. *TTC22* mRNA is a key target of *miR663A*, a miRNA that acts against inflammation during the development and progression of colon cancer [10, 11]. Therefore, we tested whether loss of function of *Ttc22* affects AOM/DSS-induced colon carcinogenesis in mice. We found that the incidence of colon cancer was similar in all of these mice treated with AOM/DSS. We detected no significant difference in the level of total RNA m6A modification in colon tissues from these mice with various *Ttc22* genotypes. This may account for the lack of difference in the prevalence of AOM/DSS induction between these mouse groups. In the current study, we cannot exclude the possibility that the doses of AOM and DSS administered were too high to display the effects of *Ttc22* inactivation. TTC22 is a scaffold protein. It is unknown where other redundant scaffold proteins may functionally compensate for defects in Ttc22 expression.

In conclusion, we report that deletion of the Ttc22 TPR4 domain disrupts Ttc22 expression and does not induce abnormal phenotypes in mice, including development, growth, or COAD induction by AOM/DSS.

## Abbreviations

COAD: colon adenocarcinoma
Ttc22: tetratricopeptide repeat domain 22
AOM: azoxymethane
DSS: dextran sodium sulfate
m6A: N6-methyladenosine
LDL: low-density lipoprotein
TCGA: The Cancer Genome Atlas
Ttc22-KO: Ttc22 knockout
SPF: specific pathogen-free
FPKM: fragments per kilobase of transcript per million mapped reads
FC: Fold Change
GO: Gene Ontology
LClllMS/MS: liquid chromatography–tandem mass spectrometry
NC: negative control
PC: positive control
DTT: dithiothreitol
SDS-PAGE: sodium dodecyl sulfate–polyacrylamide gel electrophoresis
PBS: phosphate-buffered saline
SD: standard deviation

## Funding

This work was supported by grants from the Beijing Hospital Authority Mission Plan (SML20191101) to DD and from the Science Foundation of Peking University Cancer Hospital (grant #2022-8) to TW.

## Author contributions

Conceptualization: DJD, WT, ABY; Funding acquisition: DJD; Investigation: JYL, WT, LKG, ABY, HFY, JZ; Methodology: WT; Project administration: DJD and WT; Visualization: DJD and WT; Writing – original draft: JYL, WT, and DJD; Writing – review & editing: all authors

## Competing interest

The authors declare that they have no competing interests.

## Acknowledgments

We acknowledge the support from Beijing Hospital Authority and Peking University Cancer Hospital for this project. We also extend our gratitude to staff from the laboratory animal center at Peking University Cancer Hospital for their care and maintenance of the animals.

## Notes

### Competing Interest Statement

The authors have declared no competing interest.

### Summary of Updates

We developed a rabbit polyclonal antibody against the full-length mouse Ttc22 protein, which can also bind its TPR4-deleted variant overexpressed in HEK293T cells. We detected no Ttc22 protein in the Ttc22-/- mouse colons, as determined by Western blotting with the antibody. Our new findings indicate that Ttc22 inactivation does not induce a phenotype in mice, probably because of gene redundancy or genetic robustness.

