## Supplementary material for "No phenotypes in *Ttc22* knockout mice": Figure S1

File: mouse-Ttc22-FL-(CMV-F).ab1 Run Ended: Jan 4, 2026, 6:17:44 Signal G:953 A:817 T:812 C:796
Sample: 07F_B-......419_CMV-F_TSS20260103-027-00005 Lane: 53 Base spacing: 16.60 835 bases in 12264 scans


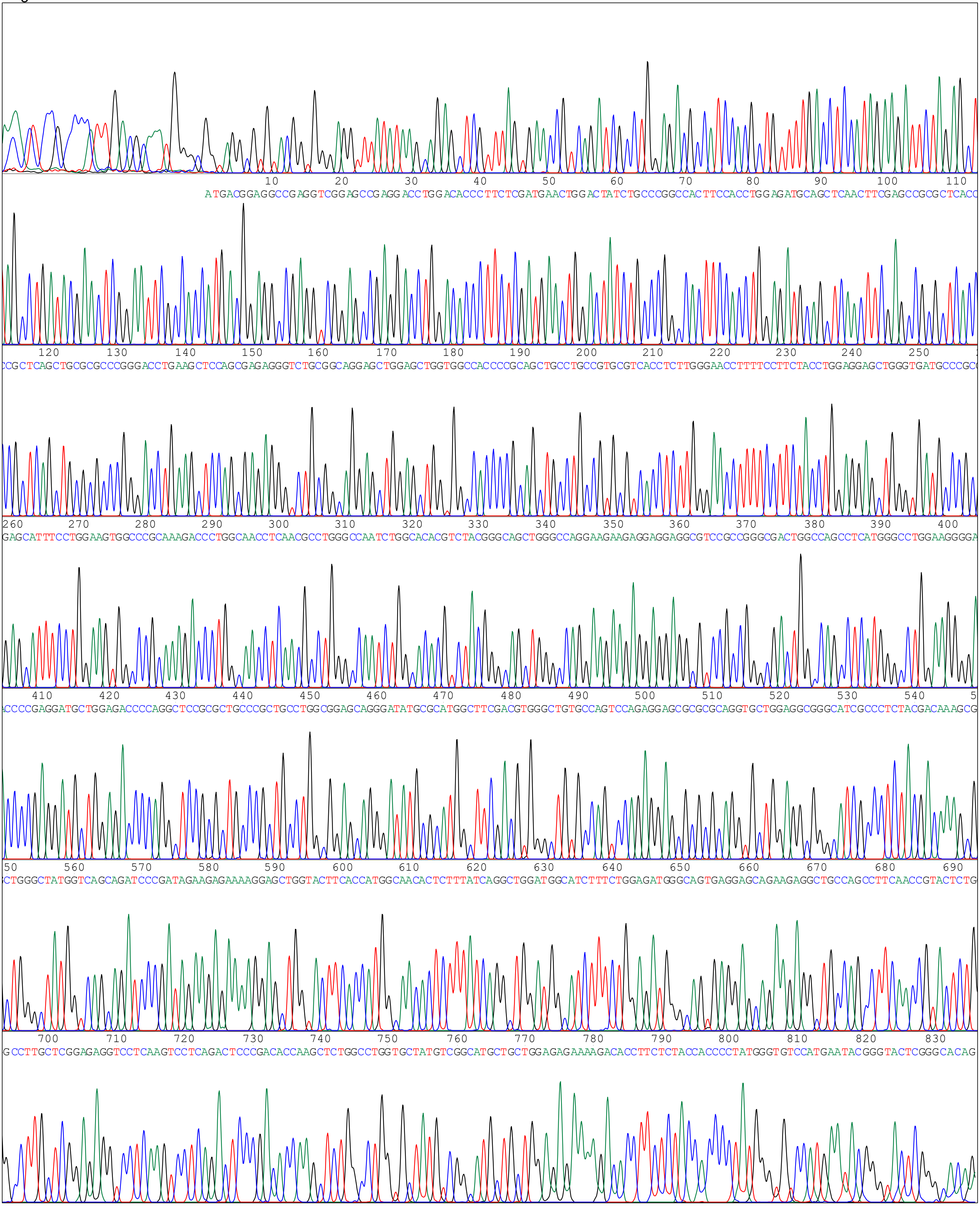
