## Supplementary material for "No phenotypes in *Ttc22* knockout mice": Figure S2

File: mouse-Ttc22-TPR1-(CMV-F).ab1 Run Ended: Dec 29, 2025, 5:27:52 Signal G:443 A:442 T:333 C:343
Sample: 06D_B-......291_CMV-F_TSS20251228-027-07917 Lane: 42 Base spacing: 16.39 898 bases in 12930 scans


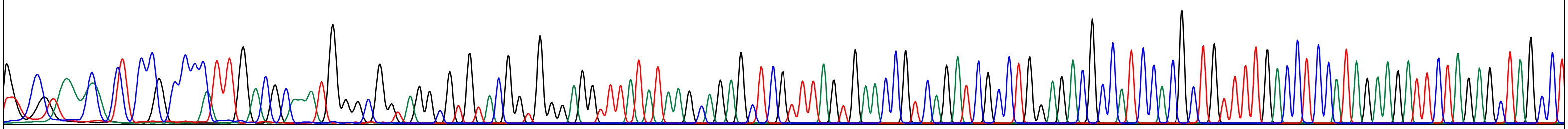


CTCAACGCCTGGGCCAATCTGGCACACGTCTACGGGCAGCTGGGCCAGGAAGAAGAGGAGGAGGCGTCCGCCGGGCGACTGGCCAGCCTCATGGGCCTGGAAGGGGACCCCGAGGATGCTGGAGAC

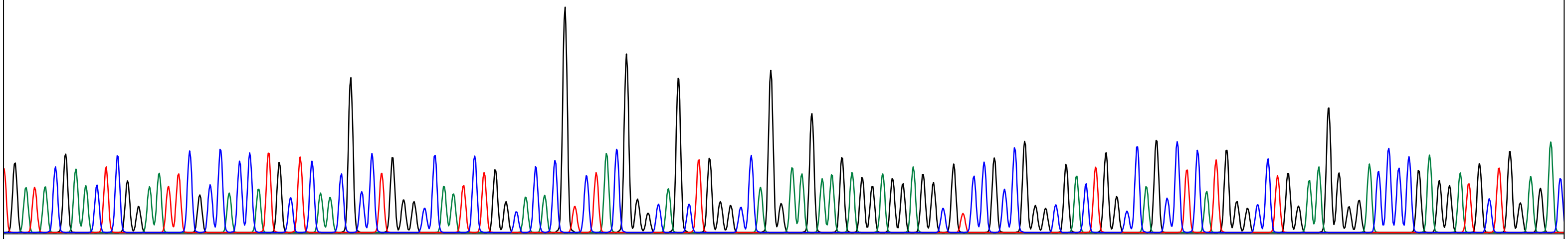


CCCCAGGCTCCGCGCTGCCCGCTGCCTGGCGGAGCAGGGATATGCGCATGGCTTCGACGTGGGCTGTGCCAGTCCAGAGGAGCGCGCGCAGGTGCTGGAGGCGGGCATCGCCCTCTACGACAAAGCGCTGGGCTATGGTCAGCAGATCCCGATAGA

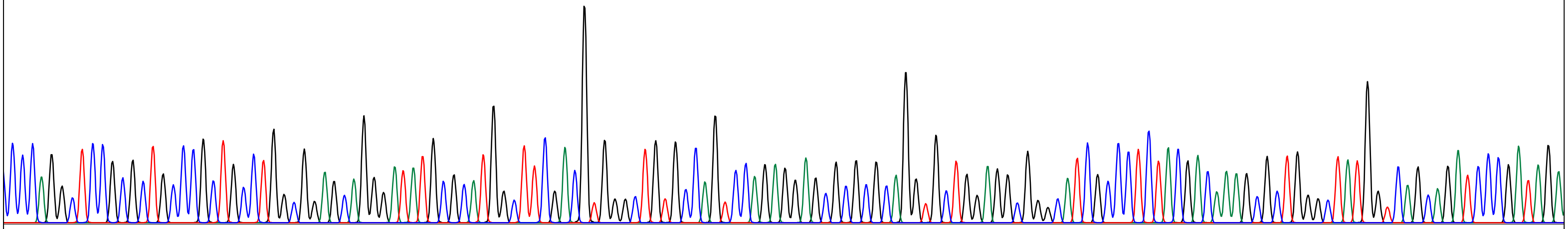


AGAGAAAAGGAGCTGGTACTTCACCATGGCAACACTCTTTATCAGGCTGGATGGCATCTTTCTGGAGATGGGCAGTGAGGAGCAGAAGAGGCTGCCAGCCTTCAACCGTACTCTGGCCTTGCTCGGAGAGGTCCTCAAGTCCTCAGACTCCCGA

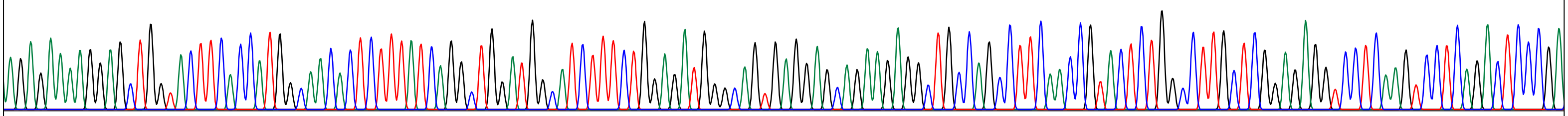


CACCAAGCTCTGGCCTGGTGCTATGTCGGCATGCTGCTGGAGAGAAAAGACACCTTCTCTACCACCCCTATGGGTGTCCATGAATACGGGTACTCGGGCACAGAGCCCCTGGACTGCTTTGGCAAGGCCATTGAGATCGCCAAGAACCAACCT

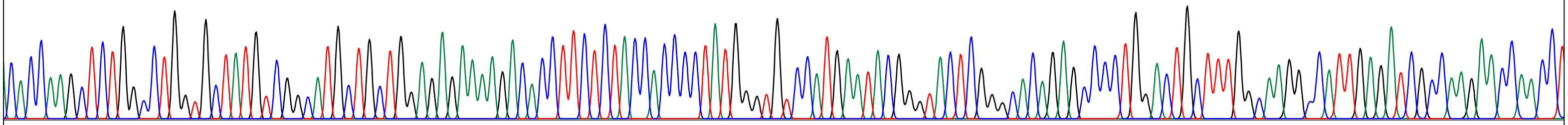


TCCCATTCTGAATCGCCTGGCCAAAATCTTCCACTTCTTGGGAAAGCAGGACATGGCCGTAGGAACCTGCAACATGGTCCTGGCGGTGCTTACAGACCCAGAGCTCAACTGGCAGGCATACTGCACCAGGGCCAAGGTCCGAATCAGAGCCTATGTC

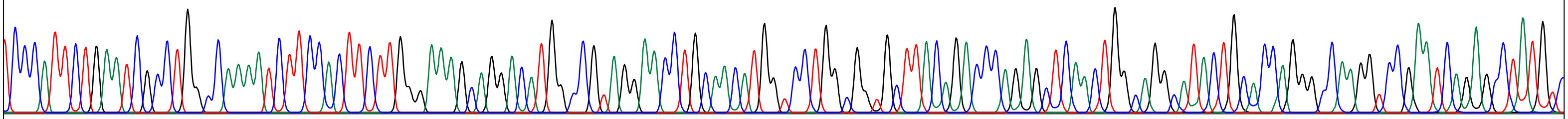


CCATGACTTGGAACGAGCCAGGGGGGGCTCGGGGTTTGCCTGACGG AACCCCTGGCCTGTGCCAAGCTGACCTGAGGAAGGGGCAAGGTGGGCCAAGCTCC GGACTACCAAAATCAGCAGGTTACACTACTGGGTGGGATGCCTGCGGAATTCTG

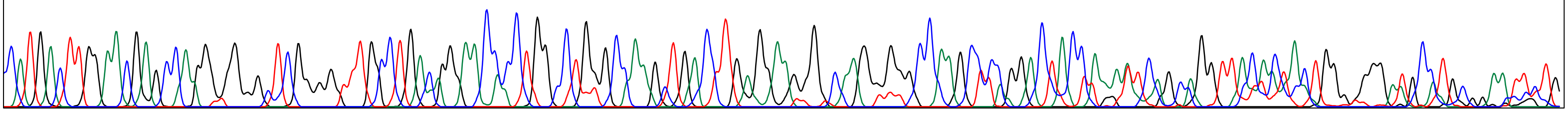
