## Supplementary material for "No phenotypes in *Ttc22* knockout mice": Figure S3

File: mouse-Ttc22-TPR4-(CMV-F).ab1
Sample: 06E_B-......15_CMV-F_TSS20251223-027-06342
Page 1 of 1

Run Ended: Dec 24, 2025, 3:28:30
Lane: 40 Base spacing: 16.38

Signal G:1753 A:2099 T:1758 C:1772
853 bases in 12535 scans

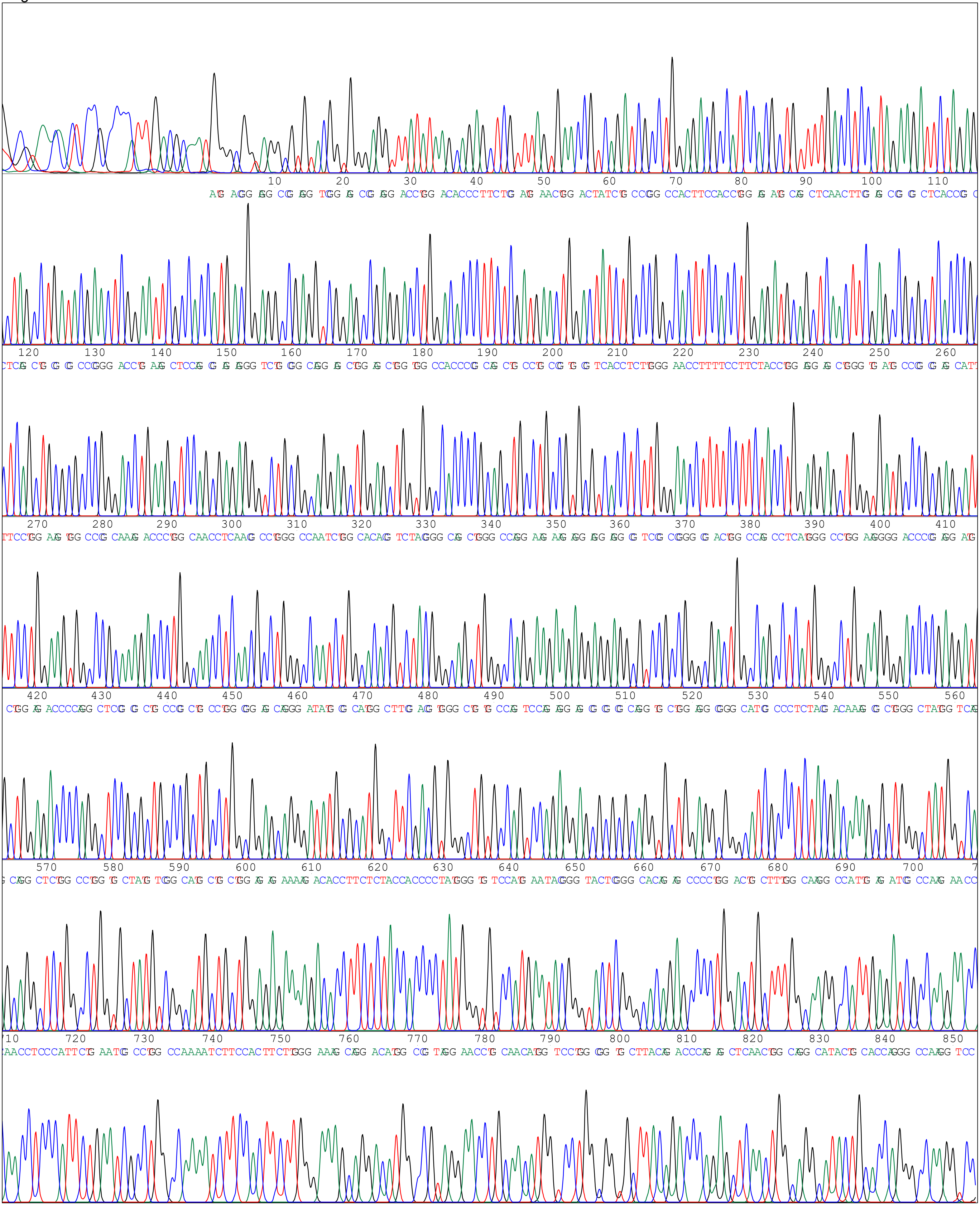
